# Peripheral immune patterns enable robust cross-platform prediction of ALS onset and progression

**DOI:** 10.1101/2025.08.26.672381

**Authors:** Luvna Dhawka, Baggio A. Evangelista, Omeed K. Arooji, Marie A. Iannone, Kyle Pellegrino, Rebecca Traub, Xiaoyan Li, Richard Bedlack, Rick B. Meeker, Todd J. Cohen, Natalie Stanley

## Abstract

Amyotrophic lateral sclerosis (ALS) progression rates vary dramatically between patients, yet the basis of this heterogeneity remains elusive, with no prognostic biomarkers existing to guide clinical decisions or stratify patients for therapeutic trials. Here, we identify a network of coordinated immune cell types, which exhibit differential disruption across progression groups. Using mass cytometry (CyTOF) to profile 2.2 million immune cells from 35 ALS patients stratified by progression rate and 9 healthy controls, we find that the extent of immune dysfunction cannot be reflected by examining differences in individual cell type frequencies. In contrast, analyses of correlation patterns between cell types revealed distinct immune organization patterns, where coordination complexity varied with disease progression. Across all progression groups, we observed striking immune reorganization in natural killer (NK) cells and a major shift from B cell/basophil coordination hubs in healthy controls to neutrophil/T cell-dominated patterns in ALS. Having established coordinated immune patterns, we developed machine learning models to further improve our ability to stratify between disease and non-disease cohorts, achieving superior performance compared to models using cell frequencies alone. Central and effector memory (CM/EM) CD4+ T cell interactions emerged as top discriminative features for disease status, while plasmacytoid dendritic cell (pDC) relationships, especially their ratio with regulatory T cells (T-regs), distinguished progression rates, supporting T-reg-based therapeutic approaches. These findings reframe ALS as a disease of immune coordination breakdown, pointing towards cell-type specific therapeutics and biomarkers that may extend beyond ALS to other neurodegenerative diseases characterized by immune dysfunction.

## Introduction

Amyotrophic lateral sclerosis (ALS) is the most prevalent motor neuron disease with an estimated prevalence of 10.5 per 100,000 persons by 2030 (Mehta et al., 2025). ALS exhibits significant clinical heterogeneity, particularly in the rate of progression from symptom onset to end-stage disease. This can range from several months to more than a decade following diagnosis (Bendotti et al., 2020). Currently, there is a lack of prognostic biomarkers that inform patient clinical progression rates. To enable improved non-invasive diagnostics, our objective is to identify cell type signatures in peripheral blood that are associated with ALS progression rates.

Multiple clinical and genetic factors have been linked with ALS progression rates, including age of onset, body mass index, C9ORF72 repeat expansions, and variations in SOD1, ATX2, and UNC13A among other genetic risk factors (Kiernan et al., 2011; Nguyen et al., 2018; Yang et al., 2019). Peripheral immune cells have emerged as an accessible source to derive ALS prognostic features based on cell frequency and phenotype. For example, it was shown that fast progressing ALS patients exhibit a reduction in CD4+FOXP3+ T regulatory cell frequency (T-regs) (Beers et al., 2017; Malaspina et al., 2015; McCauley & Baloh, 2019). In addition, increased frequencies of circulating senescent immune cells, such as late memory B cells and T-cells positive for programmed cell-death 1 (PD-1), were elevated in ALS patients compared to healthy controls (Yildiz et al., 2023). These findings are in agreement with a previous proteomics study that revealed fast-progressing ALS mononuclear cells increased markers of immune senescence (specifically senescence-associated secretory phenotype, or SASP) (Zubiri et al., 2018). More recently, increased natural killer (NK) cell frequencies were identified in ALS cohorts relative to healthy controls, and it was further shown that NK cell functional marker levels (NKG2D, NKp30, and NKp46) correlate with ALS in an age-and sex-dependent manner (Murdock et al., 2021). However, when stratified by disease progression, NK cell frequencies appeared unchanged in patients with rapidly progressing disease (Piccoli et al., 2023). Thus, distinct biomarkers of ALS progression have not been delineated, and we suspect that immune signatures may fill this void and provide prognostic utility.

In this study, we examined whether deep immunophenotyping of whole blood can generate robust immune signatures that can be used to predict ALS progression rates in a cohort of slow-progressing, standard-progressing, or fast-progressing ALS patients. We used mass cytometry (CyTOF) to generate immune signatures based on relative proportions of cell types (Stanley, 2020), and coordination patterns that enabled accurate prediction of ALS progression rate. Our immune signatures were defined based on 25 major immune populations across 2.2 million cells and revealed characteristic changes in cell type frequencies and their coordination patterns that were distinct between ALS progression groups. Key immune cell types, such as naïve CD4 T-cells and plasmacytoid dendritic cells, emerged as potential disease modifiers, biomarkers, and/or therapeutic targets. Remarkably, our CyTOF-derived immune signatures were preserved when aligned with a single-cell RNA sequencing data from an independent validation cohort (Itou et al., 2024), implying that immune networks across different platforms can be used to generate predictive models of ALS onset and progression.

## Results

### Immune profiling with mass cytometry reveals distinct immune cell populations in whole blood from ALS patients

#### Cell type identification and phenotyping

We investigated whether deep quantitative immunophenotyping of the peripheral immune system could identify robust immune signatures associated with ALS progression rates (Figure 1). We performed mass cytometry (CyTOF) analysis on whole blood samples from ALS patients (n=35) and healthy controls (n=9) (‘Discovery cohort’), to characterize immune cell populations and identify potential cell types that inform disease progression. Patients were classified into three progression groups based on the rate of change to the revised ALS functional rating scale score (ΔALSFRS-R/t): slow (n=16, ΔALSFRS-R/t < 0.5), standard (n=9, 0.5 ≤ ΔALSFRS-R/t ≤ 1.0), and fast (n=10, ΔALSFRS-R/t > 1.0) (Table 1A). Patient demographics showed similar sex distributions across groups, with a slightly younger age of onset in fast progressors (59.4 ± 10.1 years) compared to slow (65.6 ± 11.6) and standard (63.9 ± 7.5) progression groups.

**Figure 1:**
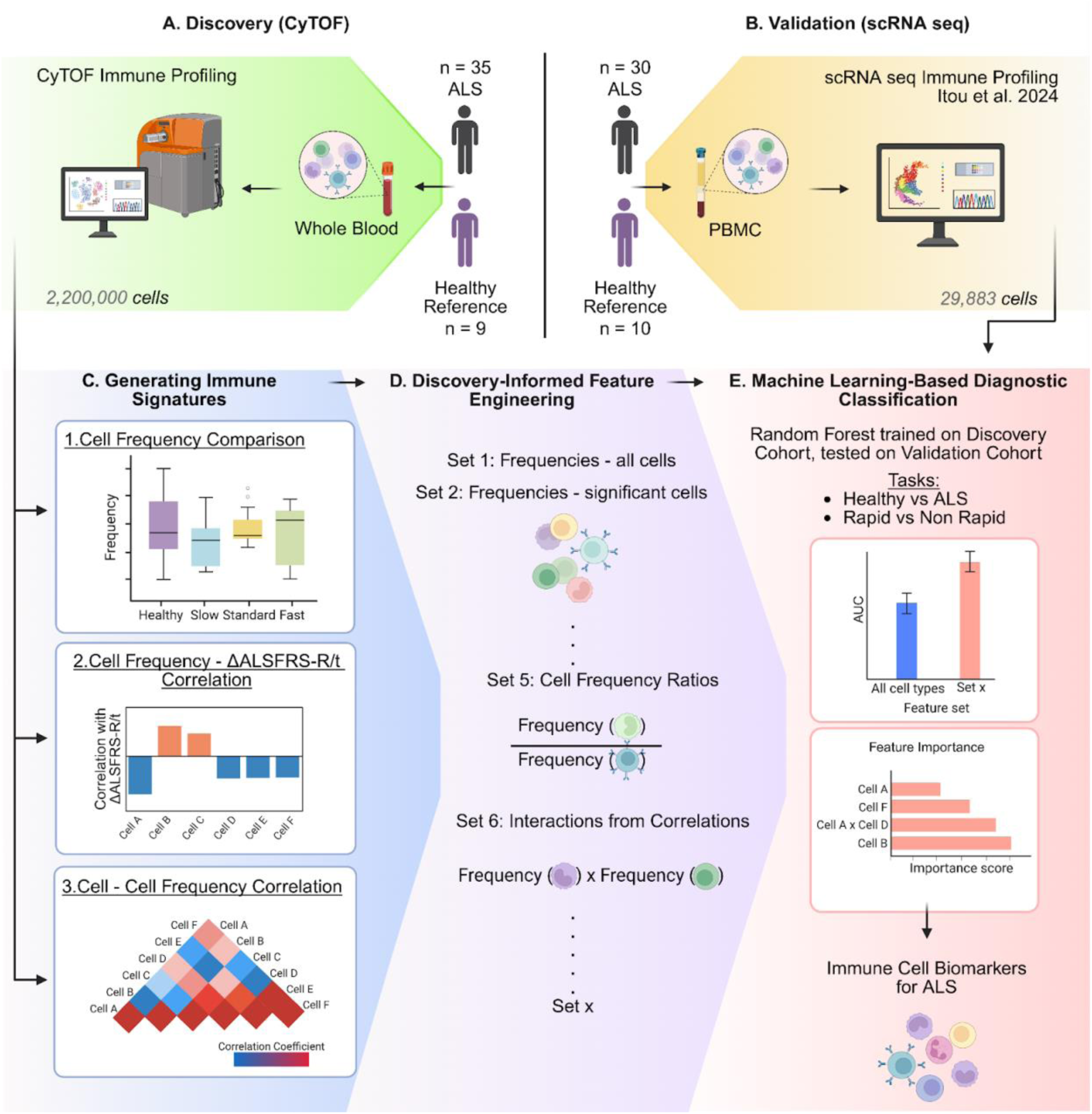
O**v**erview **of ALS Immune profiling study design and analysis.** Workflow showing the experimental design and analysis pipeline for immune profiling with CyTOF in blood from ALS donors. **(A) Discovery cohort**: Mass cytometry (CyTOF) was used to profile whole blood collected from ALS patients (n=35) across slow, standard, and fast progression groups and from healthy controls(n=9). **(B) Validation cohort:** Independent single-cell RNA sequencing (scRNA-seq) of peripheral blood mononuclear cells from ALS patients (n=30) and healthy controls (n=10) from Itou et al. 2024. The workflow proceeds through three main phases shown schematically in: **(C) Generating Immune Signatures** which involves statistical comparison of cell type frequencies across progression groups, correlation analysis between individual cell frequencies and disease progression scores (ΔALSFRS-R/t), and cell-cell frequency correlations. **(D) Discovery-Informed Feature Engineering** of signal-rich immune features based on patterns identified in **(C)**, including cell type frequencies, progression group-specific cell type frequency, cell frequency ratios, and interaction terms derived from correlation analysis (the complete feature set descriptions are provided in Supplementary Table S1). **(D) Machine Learning-Based Diagnostic Classification** trains models for ALS progression in the Discovery cohort and validates them in the Validation cohort to predict Healthy vs ALS and Rapid vs Non-Rapid progression. Feature importance analysis identifies the most informative immune features, revealing immune cell biomarkers for ALS progression.

**Table 1:**
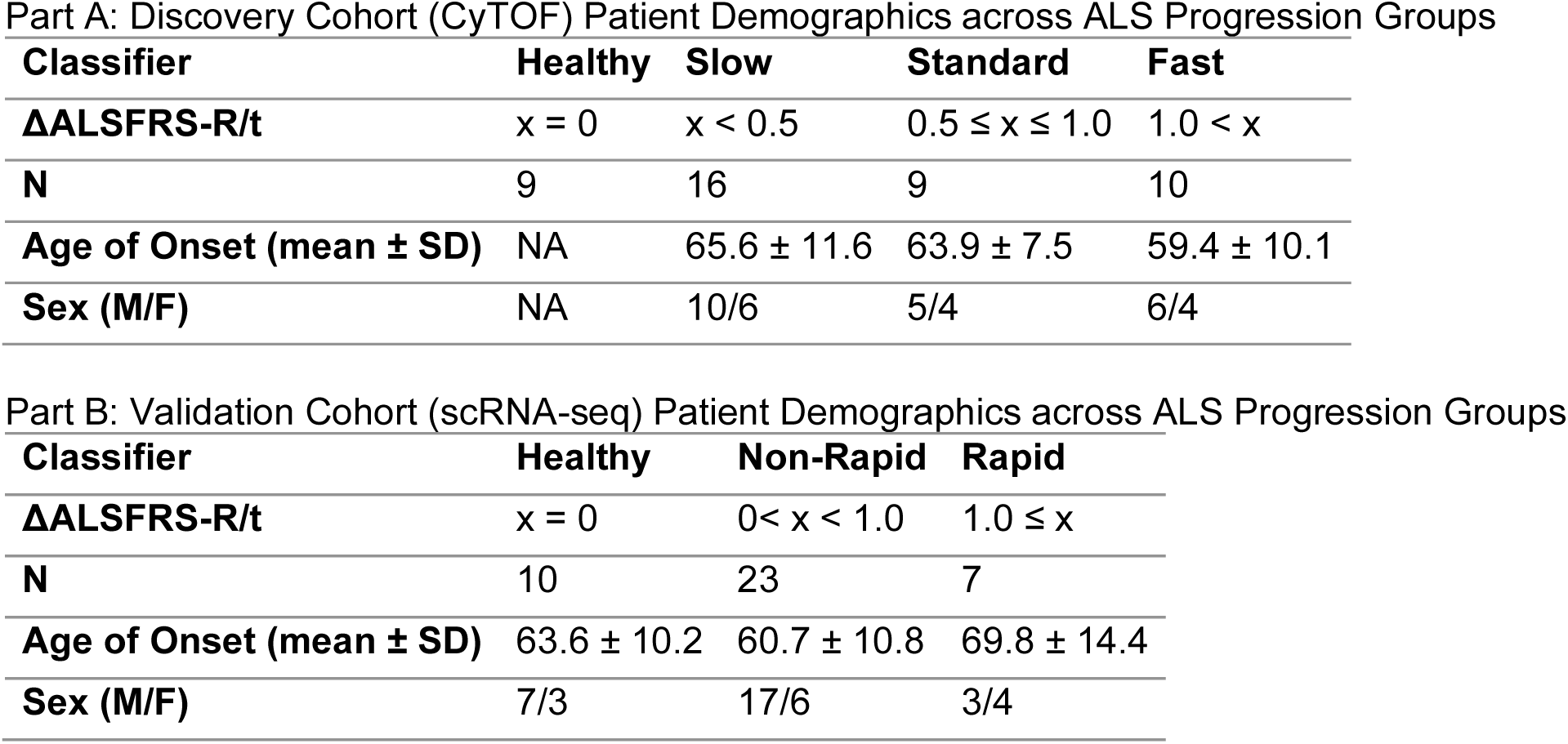

| **Classifier** | **Healthy** | **Slow** | **Standard** | **Fast** |
| --- | --- | --- | --- | --- |
| **ΔALSFRS-R/t** | $x = 0$ | $x < 0.5$ | $0.5 \leq x \leq 1.0$ | $1.0 < x$ |
| **N** | 9 | 16 | 9 | 10 |
| **Age of Onset (mean ± SD)** | NA | $65.6 \pm 11.6$ | $63.9 \pm 7.5$ | $59.4 \pm 10.1$ |
| **Sex (M/F)** | NA | 10/6 | 5/4 | 6/4 |

Part B: Validation Cohort (scRNA-seq) Patient Demographics across ALS Progression Groups
| **Classifier** | **Healthy** | **Non-Rapid** | **Rapid** |
| --- | --- | --- | --- |
| **ΔALSFRS-R/t** | $x = 0$ | $0 < x < 1.0$ | $1.0 \leq x$ |
| **N** | 10 | 23 | 7 |
| **Age of Onset (mean ± SD)** | $63.6 \pm 10.2$ | $60.7 \pm 10.8$ | $69.8 \pm 14.4$ |
| **Sex (M/F)** | 7/3 | 17/6 | 3/4 |

Using a 30-marker panel suitable for Mass Cytometry (CyTOF) [CD45, CD3, CD4, CD19, CD20, CD56, CD14, CD16, CD66b, CD123, CD11c, HLA-DR, CD294, CD45RA, CD45RO, CD27, IgD, CCR7, CD25, CD127, CD161, CD28, CD57, CD38, CXCR3, CCR6, CCR4, CXCR5, TCRγδ], we identified 25 immune cell populations from 2,200,000 cells through unsupervised Leiden clustering (Traag et al., 2019) and manual annotation based on phenotypic and functional marker expression patterns (Figure 2A-B). Our annotations captured major immune-cell lineages such as T cells (naive and memory CD4+ and CD8+ subsets, regulatory T (T-reg) cells, T helper cells, and γδ T cells), B cells (naive, memory, transitional, and plasmablasts), natural killer (NK) cells, monocytes (classical, non-classical, and transitional), plasmacytoid dendritic cells, granulocytes (neutrophils, eosinophils, basophils), and innate lymphoid cell precursors (ILCPs). As expected in whole blood, our analysis of cell-type frequencies showed neutrophils as the predominant population (64.2% of all cells), with varying proportions of other cell types that aligned with expected distributions in whole blood (Figure 2B.

**Figure 2.**
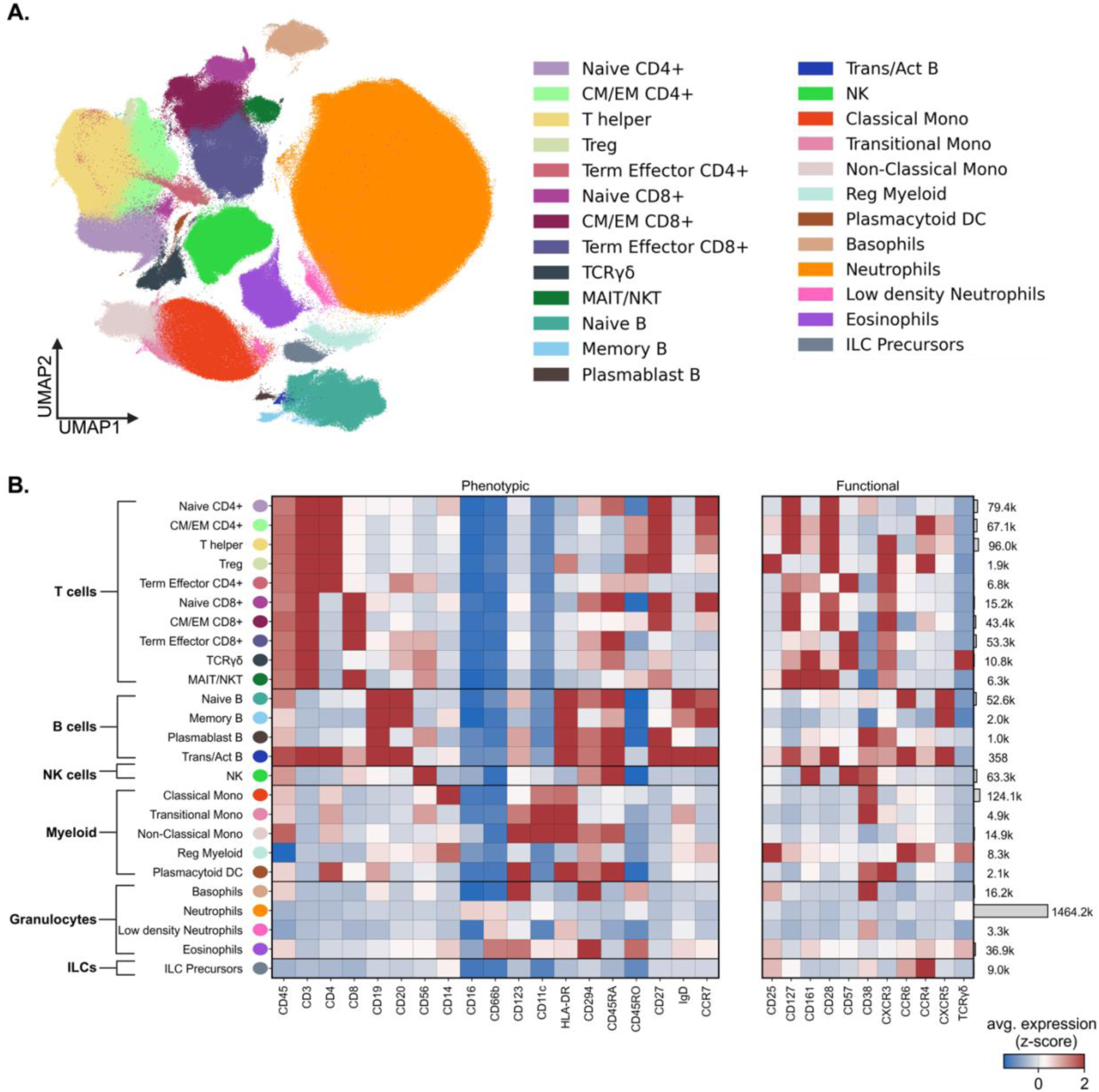
Whole blood immune cell phenotyping with CyTOF in ALS patients. **(A)** UMAP visualization of the 2,200,000 cells across all donors in the discovery cohort, colored according to the 25 identified immune cell types. **(B)** Heatmap showing average expression (z-score) of phenotypic (left) and functional (right) markers used to phenotype cell populations. The number on the right of the heatmap indicates the frequency counts of each cell type across all samples. Cell types are grouped by major lineages (T cells, B cells, NK cells, Myeloid, Granulocytes, and Innate Lymphoid Cell Precursors (ILCPs)).

### Examining differences in cell type frequencies across ALS progression groups

To identify immune cell types associated with ALS progression, we first compared cell type frequencies (i.e. abundances) across disease progression groups and healthy reference controls. Kruskal-Wallis tests revealed statistically significant differences in frequencies of many cell types, with innate lymphoid cell precursors (ILCPs), plasmacytoid dendritic cells (pDCs), and transitional monocytes showing the most prominent differences across progression groups (p=0.011, p=0.013, and p=0.034, respectively, before FDR correction) (Table 2A).

**Table 2.** Statistical analysis of cell type frequency differences across ALS progression groups. (A) Top 5 cell types showing differences across all groups by Kruskal-Wallis test. (B) Cell types with significant differences in frequency between specific groups after FDR correction (Benjamini-Hochberg procedure). (C) Cell types with statistically significant differences in frequency before but not after FDR correction. P-values were corrected for multiple comparisons within each cell type for Mann-Whitney U tests.

| **Cell Type** | **Raw p-value** | **Corrected p-value** |
| --- | --- | --- |
| Innate Lymphoid Cell Precursors | 0.011 | 0.139 |
| Plasmacytoid Dendritic cells | 0.013 | 0.139 |
| Transitional Monocytes | 0.034 | 0.248 |
| TCR$\gamma\delta$ T-cell | 0.117 | 0.585 |
| Regulatory Myeloid Cells/MDSCs | 0.133 | 0.585 |

Part B: Mann-Whitney U Tests - Significant After FDR Correction
| **Cell Type** | **Comparison** | **Raw p-value** | **Corrected p-value** |
| --- | --- | --- | --- |
| Plasmacytoid Dendritic Cells | Standard vs Slow | 0.0017 | 0.012* |
| Innate Lymphoid Cell Precursors | Healthy vs Fast | 0.0048 | 0.034* |

| Cell Type | Comparison | Raw p-value | Corrected p-value |
| --- | --- | --- | --- |
| Innate Lymphoid Cell Precursors | Fast vs Slow | 0.0190 | 0.058 |
|  | Healthy vs ALS | 0.0251 | 0.058 |
| Transitional Monocytes | Standard vs Slow | 0.0188 | 0.088 |
|  | Fast vs Slow | 0.0251 | 0.088 |
| Regulatory Myeloid Cells/MDSCs | Healthy vs ALS | 0.0215 | 0.103 |
|  | Healthy vs Slow | 0.0293 | 0.103 |
| Plasmacytoid Dendritic cells | Fast vs Slow | 0.0219 | 0.077 |
| TCR$\gamma\delta$ T-cell | Fast vs Slow | 0.0424 | 0.178 |
| Regulatory T cells | Healthy vs Fast | 0.0455 | 0.219 |

*Note: \* indicates significance after FDR correction ( $p < 0.05$ ). MDSCs = Myeloid-derived Suppressor Cells. All p-values were corrected for multiple comparisons using the Benjamini-Hochberg procedure, applied separately to Kruskal-Wallis tests and within each cell type for Mann-Whitney U tests.*

Pairwise comparisons between progression groups using Mann-Whitney U tests uncovered two significant differences after FDR correction: pDCs were less abundant in standard progressors than in slow progressors (FDR-corrected p=0.012), and, ILCPs had significantly lower frequency in fast progressors than in healthy controls (FDR-corrected p=0.034) (Figure 3, Table 2B).

**Figure 3.**
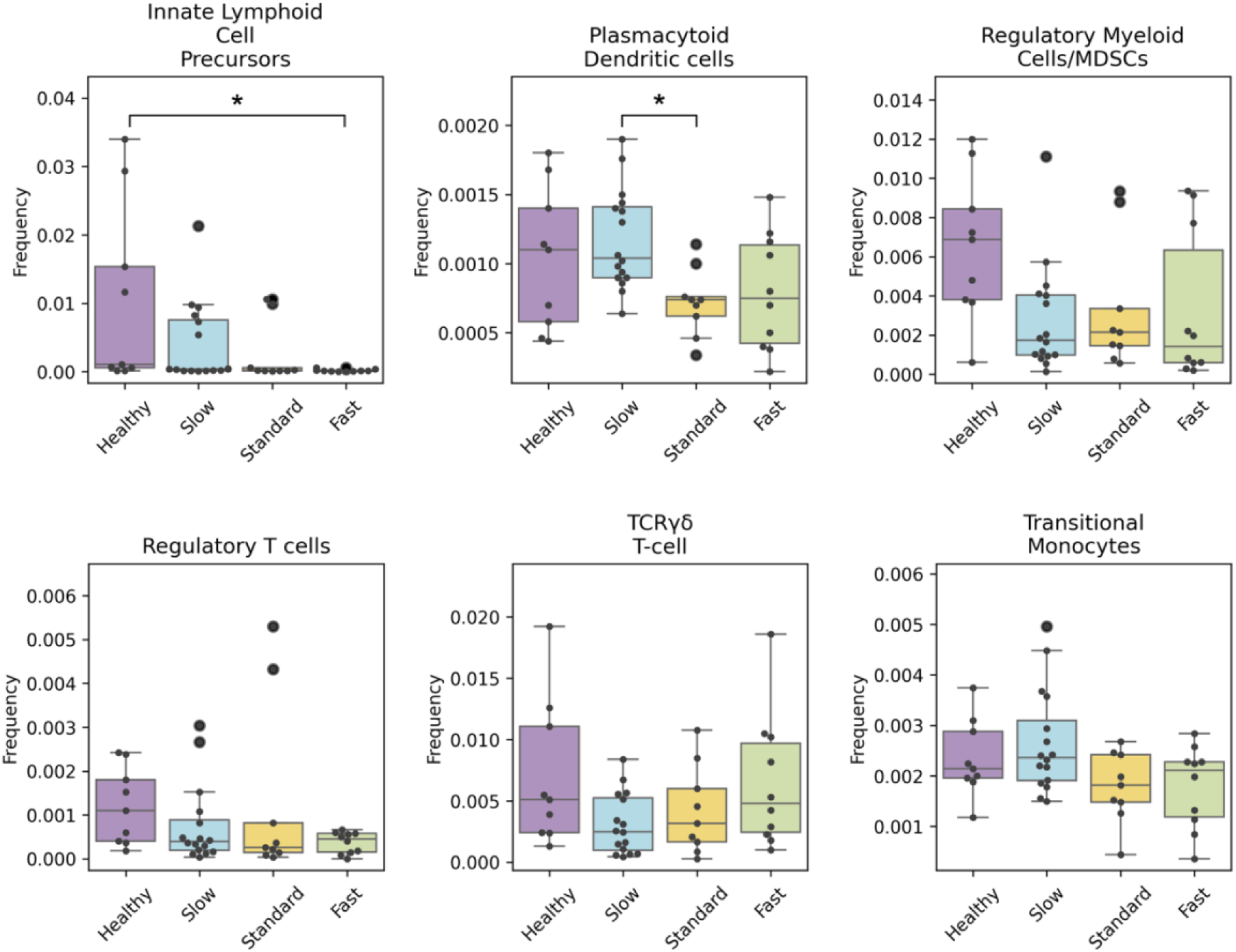
Cell type frequencies across ALS progression groups. Box plots show frequencies of selected immune cell populations with prominent differences across donors, grouped by progression group. Cell types visualized include Innate Lymphoid Cell Precursors, Plasmacytoid Dendritic cells, Regulatory Myeloid Cells/MDSCs, Regulatory T cells, TCRγδ T-cells, and Transitional Monocytes. Asterisks indicate statistical significance (Mann-Whitney U test, FDR corrected, *p < 0.05). Groups are colored according to progression category: Healthy (purple), Slow (blue), Standard (yellow), and Fast (green).

Beyond these two observations, we noted several distinct patterns in cell frequencies across healthy reference controls and the ALS progression spectrum (Figure 3, Figure S1). First, we noted a progressive enrichment in some cell types across slow, standard, and fast progression states compared to healthy reference controls, which included naïve B cells and naïve CD4 T cells (neither were statistically significant). Second, we noted a progressive depletion in some cell types across slow, standard, and fast progression states compared to healthy reference controls, including ILCPs (Healthy > ALS, not statistically significant after FDR correction; Healthy > Fast, FDR-corrected p=0.034; Slow > Fast, not statistically significant after FDR correction) and NK cells (not statistically significant) show this trend. Third, some cell types showed similar frequencies across all ALS progression groups that differed from healthy reference controls. Regulatory myeloid cells/MDSCs (Healthy > ALS, not statistically significant after FDR correction; Healthy > Slow, not statistically significant after FDR correction) and T-reg cells (Healthy > Fast, not statistically significant after FDR correction) follow this pattern. Finally, some cell types were differentially abundant in specific ALS progression groups compared to other progression groups. These include pDCs (Slow > Standard, FDR-corrected p=0.012; Slow > Fast, not statistically significant after FDR correction), transitional monocytes (Slow > Standard, not statistically significant after FDR correction; Slow > Fast, not statistically significant after FDR correction) and TCRγδ T-cells (Fast > Slow, not statistically significant after FDR correction), whose frequency differences suggest potential immunological signatures unique to each progression group.

In addition to these clear differences in cell type frequencies among progression groups, several other cell types showed more variable frequencies that could not be distinguished among cohorts, including memory CD8+ T cells, terminal effector T cells, MAIT/NKT-like cells, classical monocytes, and T helper cells (Supplementary Figure S1).

### **C**orrelation analysis identifies immune cell types associated with disease progression

To further investigate relationships between immune cell type frequencies and disease progression, we examined how individual cell type frequencies correlate with ΔALSFRS-R/t scores using Spearman’s correlation on log-transformed cell-type frequencies. We assessed these correlations in two contexts. We first examined correlations while including healthy controls (ΔALSFRS-R/t = 0). We also computed correlations with the exclusion of healthy controls to distinguish relationships spanning the entire healthy-to-ALS progression spectrum compared to those that may be specific to ALS progression states (Figure 4 A-B).

**Figure 4.**
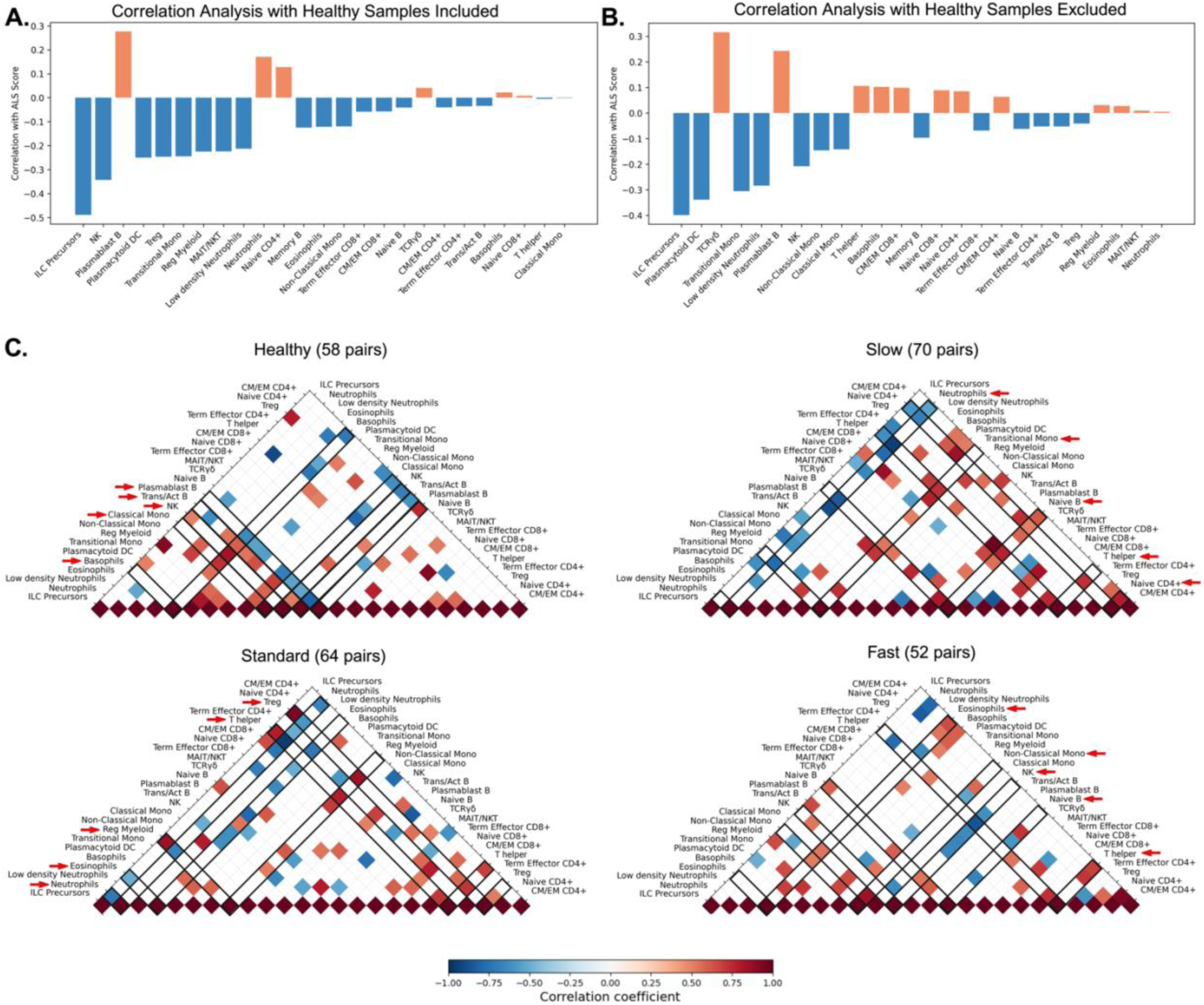
Correlation patterns between cell type frequencies in each progression group. (A-B) Bar plots show Spearman correlation coefficients between cell type frequencies and ALSFRS-R/t scores, with **(A, left)** and without **(B, right)** healthy controls included in the analysis. Negative correlations (blue) indicate cell types that decrease with faster disease progression and positive correlations (orange) show cell types that increase with faster disease progression. **(C)** Correlation matrices show inter-cell type frequency relationships across the four groups (healthy, slow, standard, fast). Red indicates positive correlations; blue indicates negative correlations as indicated by the correlation coefficient color bar. Red arrows indicate the top 5 cell type *hubs* (e.g. cell types with the highest number of significant frequency correlations with other cell types) in each group, with black lines outlining all the significant correlations for each identified hub cell type across the correlation matrix. Only correlations with |r| ≥ 0.5 and p < 0.001 (FDR corrected) are shown.

With healthy controls included, ILCPs showed the strongest negative correlation with progression rate (r=-0.488, FDR-corrected p=0.0192), indicating lower frequencies are associated with faster disease progression (Figure 4A). NK cells also displayed a negative correlation (r=-0.343, p=0.023, FDR-corrected p = 0.286), while several cell types including plasmablast B cells showed positive correlations (r=0.276, p=0.070, FDR-corrected p=0.452), indicating higher frequencies with faster progression (Figure 4B). These findings align with the frequency patterns observed in Figure 3, with respect to the progressive depletion of innate immune components across the healthy-ALS progression spectrum.

When examining correlations in cell type frequencies in only ALS patients, a slightly different correlation pattern emerged. ILCPs were still the most strongly negatively correlated cell type with ΔALSFRS-R/t scores (r=-0.399, p=0.018, FDR-corrected p = 0.444), indicating that their frequencies are also useful for delineating ALS progression groups. Instead of NK cells, pDCs emerged with the second strongest negative correlation (r=-0.338, p=0.047, FDR-corrected p = 0.465), while TCRγδ T-cells exhibited the strongest positive correlation (r=0.315, p=0.065, FDR-corrected p= 0.465) with progression rates (Figure 4B).

From the above correlation analyses among healthy and ALS cohorts, we conclude the following: (1) correlation between individual cell types and disease progression were modest at best, with no single cell type showing strong independent correlation with ALS progression; (2) some cell types, such as NK cells, display abundances that more strongly differentiate healthy from ALS as opposed to differentiating between ALS progression states; and (3) other cell types, such as TCRγδ T-cells, have frequencies that appear to be more associated with ALS progression rate.

### Insights from progression-specific inter-cell type correlation patterns

To understand how coordinated relationships (i.e. whole network changes) between immune cell types might change with disease progression, we analyzed correlations between cell types within each progression group.

We analyzed how correlations between cell types might collectively infer regulatory relationships where the number and strength of a cell type’s correlation with other cell types suggest its role in immune coordination. Strong positive correlations (where two cell populations have frequencies, which increase or decrease in tandem) may imply coordinated or shared regulatory mechanisms, while negative correlations might reflect inhibitory interactions or opposing responses to disease-related environments. The number of significant correlations that a cell type maintains (referred to as connectivity) can indicate its potential importance as an immune coordination hub or *cell hub*, in different disease states. Using stringent criteria (|r| ≥ 0.5, p < 0.001 after FDR correction, described in methods), we identified a large set of significant correlations between cell types across healthy controls and ALS progression groups (Figure 4C).

#### Coordination complexity varies across ALS progression groups with highest coordination in slow progressors

The correlation matrices in Figure 4C show distinct coordination patterns and densities between immune cell types across progression groups, with the most striking observation being differences in the overall density of coordination connectivity between progression groups. Slow progressors displayed the most complex coordination with 70 significant correlation pairs, followed by standard progressors (64 pairs), healthy controls (58 pairs), and fast progressors showing the least connectivity (52 pairs).

#### Immune cell hubs shift across ALS progression groups

Assessing cell type connectivity revealed striking shifts in hub cells across progression groups (Figure 4C, Supplementary Figure S2).

In healthy controls, B cell populations were inferred to be important coordination hubs, with transitional/activated B cells (10 connections), plasmablast B cells (9 connections), and basophils (8 connections) showing the highest connectivity. NK cells and classical monocytes also exhibited many connections (7 connections each).

When comparing ALS progression states to the healthy controls, we observed distinct reorganization patterns in each group. Slow progressors showed a reorganization of the primary cell hubs. Neutrophils and T helper cells (11 connections each) were the primary coordination hubs, with naive B cells and naive CD4+ T cells (10 connections each) and transitional monocytes (8 connections) also showing high connectivity. Standard progression was also characterized by distinct coordination reorganization compared to healthy controls, with eosinophils and neutrophils (9 connections each) showing up as main hubs, followed by T helper cells, regulatory myeloid cells, and T-regs (8 connections each). Finally, fast progressors displayed another distinct pattern of immune coordination when compared to healthy controls, with T helper cells, naive B cells, and NK cells (7 connections each) serving as primary hubs, followed by non-classical monocytes and eosinophils (6 connections each).

#### Immune coordination complexity reveals distinct architectures across ALS progression groups

Beyond shifts in individual cell hubs, we observed fundamental changes in the underlying architecture of innate and adaptive coordination in the immune system, categorized in terms of innate-adaptive crosstalk, adaptive immune coordination, and innate immune coordination (Supplementary Figure S3).

Healthy controls exhibited prominent cross-talk between the innate and adaptive components of the immune system, given that 53% of their statistically significant cell correlation pairs were between innate and adaptive components (31/58 pairs). Otherwise, 22% of correlated pairs (13/58) were within the adaptive immune system and 24% (14/58 pairs) were in the innate immune system. This baseline pattern shifted distinctly across ALS progression groups. Slow progressors showed 31% of significant cell-cell correlations in the adaptive immune system, in contrast to only 16% of coordination patterns in innate cell types. Standard progressors exhibited 33% cell coordination patterns in adaptive immune components and 16% innate immune coordination. Fast progressors displayed 27% of significant coordination patterns in adaptive immune components and 23% innate immune coordination, with proportional coordination patterns most similar to healthy controls across all three categories.

The proportion of innate-adaptive crosstalk remained relatively stable across all groups (52% in slow, 52% in standard, and 51% in fast progressors).

#### Unique and conserved inter-cell type correlation signatures highlight disease progression-specific immune coordination

We next compared which significant correlations were unique or shared across progression groups (Figure 5A). For each progression group and shared ALS set, we highlighted the top 5 correlation pairs based on correlation coefficient and p-values (from a much larger set of unique correlations), with positive correlations labeled in red and negative correlations in blue.

**Figure 5.**
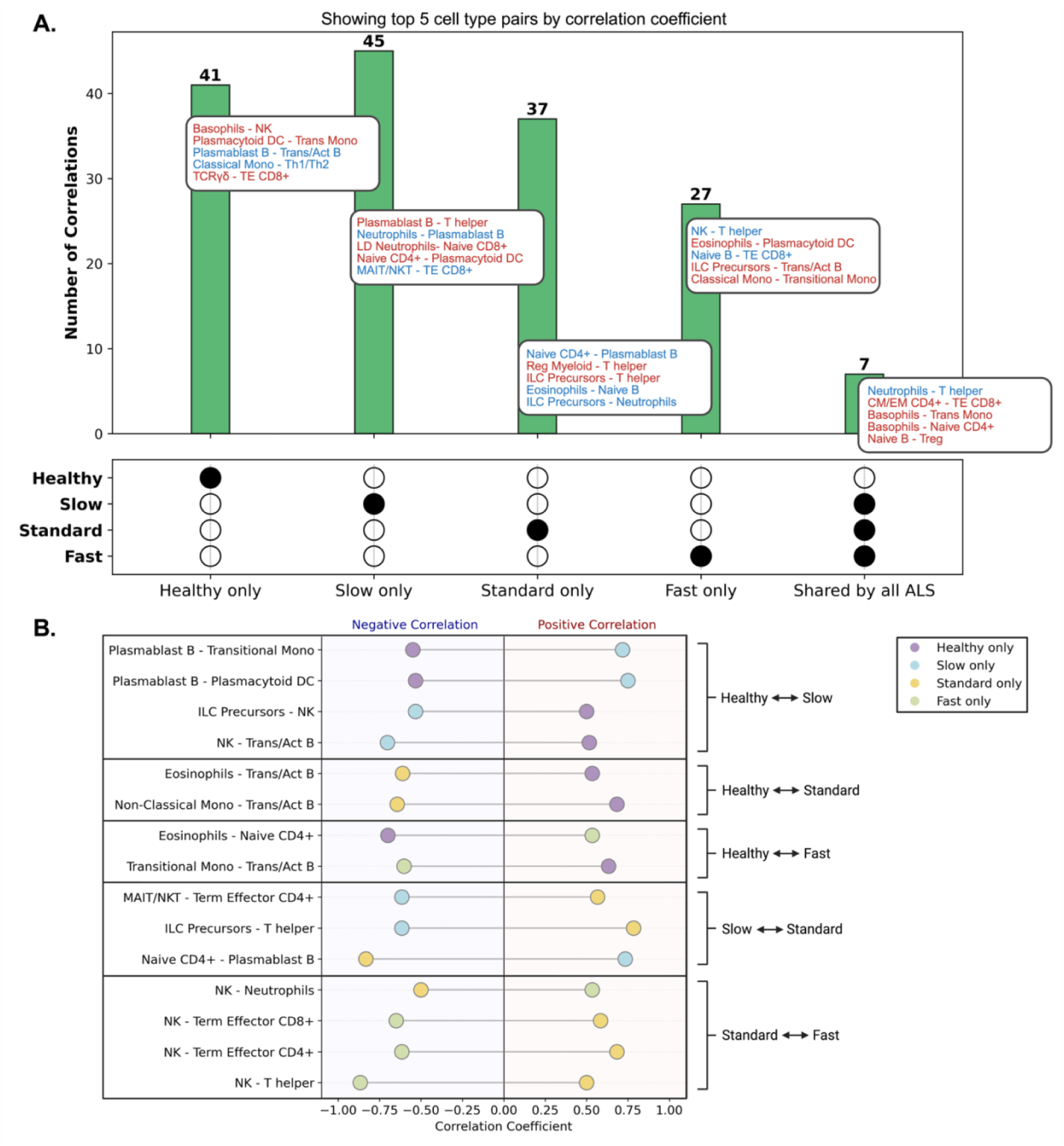
Immunological correlation coordination patterns are specific to progression groups. **(A)** UpSet plot shows the number and distribution of significant correlations between cell types based on their frequencies (|r| ≥ 0.5, p < 0.001) across progression groups. Filled circles below each bar indicate which progression group(s) the pairs of cell type correlations are significant in. Text boxes list the top five cell type pairs (based on Spearman correlation coefficient) contributing to each category. Red labels are for positive correlations; blue labels are for negative correlations. **(B)** A visualization of the correlations between cell type frequencies that switch correlation patterns between specific pairs of progression groups. Only correlations included in the UpSet plot in **(A)** are shown. Points indicate the correlation coefficient between the indicated pair of cell type frequencies in the indicated progression groups (progression groups indicated by color).

Our analysis revealed core immune coordination changes shared across ALS progression groups. Seven correlation patterns were conserved across all ALS progression groups while absent in healthy controls, representing potential disease-associated immune signatures. Most striking was the strong negative coordination between neutrophils and T helper cells (r=-0.93). This was accompanied by naive B cells coordinating with both T-regs (r=0.58) and regulatory myeloid cells/MDSCs (r=0.53). Additionally, basophils showed increased connectivity with naive CD4+ T cells (r=0.62) and transitional monocytes (r=0.70).

Beyond these shared patterns, we observed distinct coordination complexity and reorganization across progression groups. Healthy controls maintained robust innate immune coordination among their 41 unique correlation pairs, as exemplified by strong basophil interactions with NK cells (r=0.83) and classical monocytes (r=0.73), alongside pDC coordination with transitional monocytes (r=0.82). In contrast, ALS progression groups showed different coordination architectures with varying complexity: slow progressors (45 unique pairs), standard progressors (37 pairs), and fast progressors (27 pairs).

Slow progressors displayed extensive neutrophil-lymphocyte interactions, including strong negative regulation between neutrophils and plasmablast B cells (r=-0.88). pDCs showed distinct coordination patterns across progression groups: with transitional monocytes in healthy controls (r=0.82), shifting to naive CD4+ T cells in slow progressors (r=0.85), and connecting with eosinophils in fast progressors (r=0.68). Fast progressors, despite having the fewest unique correlations, featured a distinctive signature: connections between ILC precursors and transitional B cells (r=0.66).

Only two correlation patterns were preserved across all groups, including healthy controls, with terminal effector CD4+ and CD8+ T cell coordination showing progressive weakening from healthy controls (r=0.92) through slow (r=0.83) and standard (r=0.78) to fast progressors (r=0.70).

Additionally, we observed fundamental alterations in the direction of immune cell relationships across progression groups (Figure 5B). NK cells showed the most frequent sign-switching interactions, including reversals with T helper cells (positive in standard progressors, r=0.50, but strongly negative in fast progressors, r=-0.87) and with terminal effector T cells (negative in fast progressors but positive in standard progressors). pDCs also demonstrated extensive coordination pattern reorganization, with their relationships switching direction across multiple progression groups. ILCPs, the cell type most strongly associated with disease progression in our dataset, also showed sign reversals, which are described as switching from positive correlations with NK cells in healthy controls (r=0.50) to negative correlations in slow progressors (r=-0.56). When comparing each ALS progression group to healthy controls, we observed distinct patterns of coordination reorganization, with slow progressors showing shifts toward neutrophil-T cell coordination, standard progressors developing granulocyte-dominated coordination, and fast progressors maintaining mixed coordination but with reduced overall connectivity.

### Machine learning models trained on immune signatures can classify patients according to ALS status and progression

We next investigated whether immune cell frequencies derived from our CyTOF analysis could serve as predictive biomarkers for ALS status and progression rate. We trained models using our CyTOF dataset and validated them on a publicly available single-cell RNA sequencing dataset of peripheral blood mononuclear cells (PBMCs) from ALS patients and healthy controls (Itou et al. 2024). We sought to demonstrate this cross-platform validation as an indicator of the robustness of immune signatures across different technologies and independent cohorts.

The scRNA-seq validation cohort included 40 individuals (30 ALS patients and 10 healthy controls), with ALS patients classified as either Rapid (ΔALSFRS-R/t > 1.0, n=7) or Non-Rapid (ΔALSFRS-R/t ≤ 1.0, n=23) progressors (Table 1B). To align with the scRNA-seq dataset’s classification scheme, we re-classified our “Slow” and “Standard” groups to “Non-Rapid,” and our “Fast” group to “Rapid.” We developed multiple distinct feature sets to capture different aspects of immune dysregulation, including basic cell type frequencies, cell type ratios, and interaction terms derived from correlation patterns (Table 3, Supplementary Table S1). As different feature sets capture different aspects of immune coordination, testing multiple feature sets allowed us to identify which immune relationships were most predictive of disease status and progression.

**Table 3:** Selected feature sets used for classification of ALS progression (full list in supplementary table 1)

| Feature Set Name | Description |
| --- | --- |
| All cell types | Frequencies of all cell types |
| Significant cell types with ratios | Significant cell type frequencies plus ratios between significant cells |
| Significant cell types with task-specific correlation interactions | Significant cell type frequencies plus task-specific correlation interactions |

*Note: All cell type frequencies were log-transformed before analysis to improve distribution normality and model performance.*

Our Random Forest models successfully classified both disease presence (Healthy vs. ALS) and progression rate (Non-Rapid vs. Rapid) in the independent scRNA-seq validation cohort (Figure 6B and 6C, left panels). Importantly, feature sets incorporating cell type relationships significantly outperformed basic frequency features for both classification tasks. For Healthy vs. ALS classification, Feature Set 20 (Significant cell types with task-specific correlation interactions; full description in Supplementary Table S1) achieved significantly higher performance than using Feature Set 1 (All cell types) (Figure 6B left, AUC 0.83 vs. 0.50, p = 0.00254). Similarly, for Non-Rapid vs. Rapid classification, ‘Significant celltypes with ratios’ showed improved performance over the baseline feature set (Figure 6C left, AUC 0.65 vs. 0.51, p = 0.00656).

**Figure 6.**
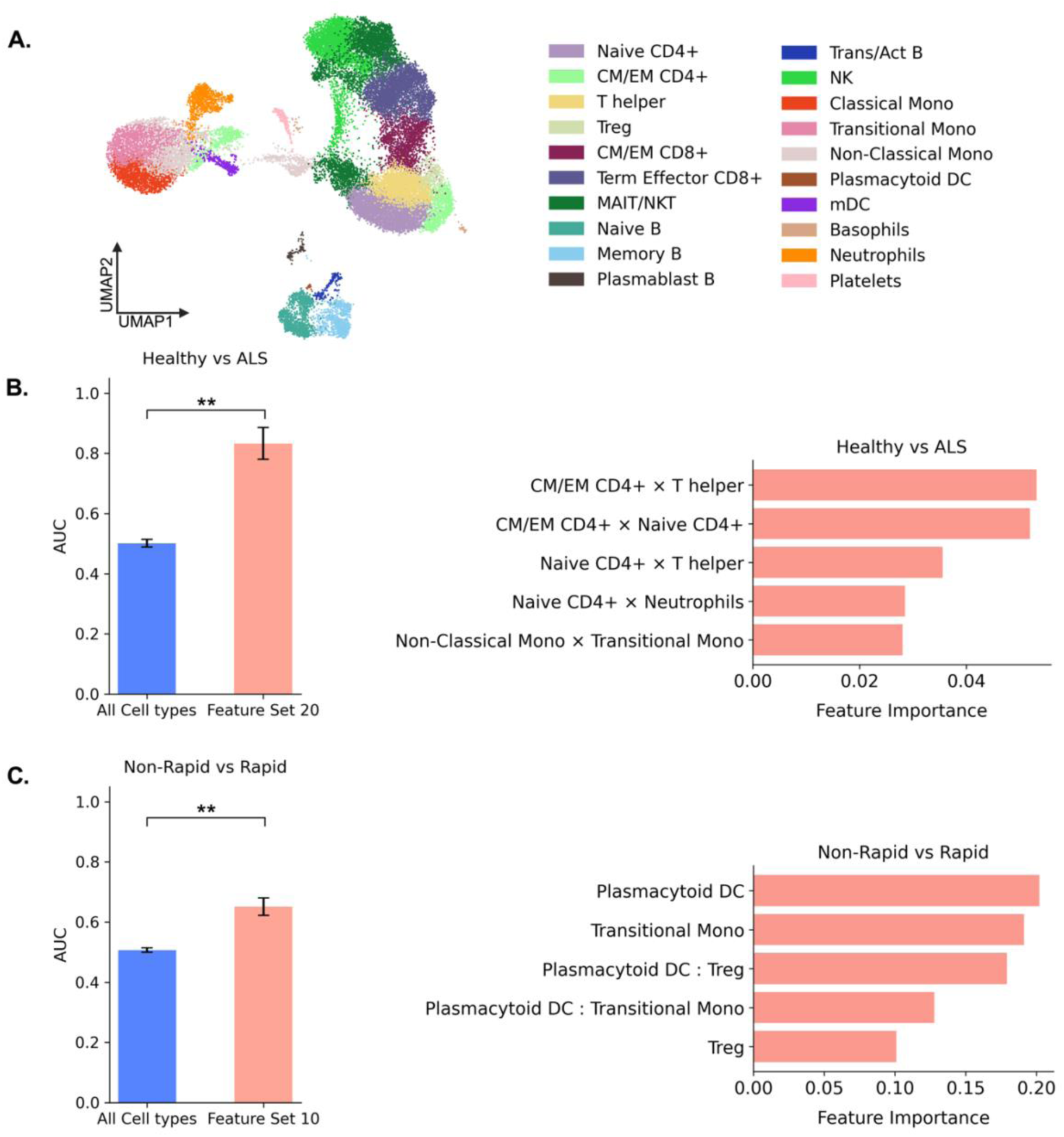
Validation of CyTOF-derived immune features using scRNA-seq profiles of PBMCs in an independent ALS cohort. **(A)** UMAP visualization of 20 immune cells profiled with scRNA-seq, colored by cell type. **(B)&(C)** Left panels show the classification accuracy between the baseline feature set (Feature Set 1: All Cell types) and the best feature sets for each task. Bar charts show the AUC averaged across 5 independent runs with standard error bars. The right panels show the feature importance plots with the 5 most important features for each classification task using the best feature sets, determined by random forest built-in feature importance scores. **(B)** For classifying healthy and ALS donors, Feature Set 20 (significant cell types with task-specific correlation interactions; see Supplementary Table S1) showed significantly higher performance than All Cell types (p = 0.00254, **). Interaction terms (×) between T cell subsets were most informative. **(C)** For Non-Rapid vs Rapid, Feature Set 10 (significant cell types with ratios; see Supplementary Table S1) showed improved performance over All Cell types (p = 0.00656, **). Plasmacytoid dendritic cells and their ratios (:) with other cell types were the most important features, consistent with their significant reduction in standard versus slow progressors.

Feature importance analysis revealed distinct patterns for each classification task. For Healthy vs. ALS discrimination, interaction terms between T cell subsets emerged as the most informative features, with central and effector memory (CM/EM) CD4+ × T helper cells and CM/EM CD4+ × naive CD4+ interactions among the top contributors (Figure 6B right). For Non-Rapid vs. Rapid classification, pDCs and their relationships with T-regs and transitional monocytes were consistently identified as key features (Figure 6C right), aligning with our correlation coordination analysis showing altered relationships between these cell types across progression groups.

## Discussion

This is the first study, to the best of knowledge, to systematically demonstrate that immune signatures derived from cell coordination patterns provide superior markers of disease progression compared to individual cell type frequencies alone. Statistical analysis of frequency differences of immune cell types between progression groups revealed statistically significant frequency differences between ALS progression groups only in plasmacytoid dendritic cells and innate lymphoid cells after correction for multiple testing. However, when examining differential correlation patterns between cell types, we uncovered extensive immune system reorganization, with 52-70 statistically significant cell-cell relationships per group. We developed immune signatures based on these coordination patterns, which dramatically outperformed individual cell frequencies in cross-platform validation (Healthy vs ALS: AUC of 0.83 vs. AUC of 0.50 with frequency features only, p = 0.00254; Rapid vs Non-Rapid: AUC of 0.65 vs. AUC of 0.51, p = 0.00656). Our results suggest that neuroinflammation, which has been implicated in ALS progression (Murdock et al., 2025), is characterized by unique immune coordination patterns (i.e. network changes) rather than by changes in individual cell type abundances. These findings are in line with reported immune cell interactions in neurodegenerative conditions, including microglia-T cell interactions in tauopathy (X. Chen et al., 2023; Mason et al., 2025), and CXCR6+ CD8+ T cell-microglia coordination in Alzheimer’s disease (Su et al., 2023).

Collectively, our results reveal tangible leads and targetable ALS-specific patterns of immune dysfunction with potential for diagnostic utility.

The observed changes in cell type frequencies suggest immune reorganization in ALS. First, innate lymphoid cell precursors (ILCPs) showed the strongest correlation with disease progression (r=-0.488, FDR-corrected p=0.0192); their frequencies decreased with increasing ΔALSFRS-R/t scores. ILCPs give rise to innate lymphoid cells (ILCs), such as group 1 ILCs or natural killer cells (Klose & Artis, 2020), which can directly kill motor neurons in cell models of ALS (Garofalo et al., 2020). Our analysis supports recent studies (Murdock et al., 2021; Piccoli et al., 2023), which revealed no significant correlation between total NK cell frequency and ALS progression rate, though it remains of interest to investigate the relationship between ILCP differentiation and neurotoxic NK cell development in ALS progression groups. Second, plasmacytoid dendritic cells (pDCs) are significantly reduced in standard compared to slow progressors (FDR-corrected p=0.012). pDCs serve as specialized bridges between innate and adaptive immunity through their capacity to produce type I interferons and present antigens to T cells (Colonna et al., 2004; Guery & Hugues, 2013). Their context-dependent function as either tolerogenic or immunogenic cells (Matta et al., 2010) may contribute to the disrupted coordination patterns observed across progression groups. Third, regulatory T cells (T-reg) displayed similar depletion patterns in fast progressors compared to healthy controls. These findings are consistent with T-reg dysfunction in rapidly progressing ALS patients (Beers et al., 2017; Henkel et al., 2013; Sheean et al., 2018; Yazdani et al., 2022).

Beyond these individual population changes, we also examined how more complex coordinated relationships between different immune cell types emerged and varied across progression groups. First, the number of statistically significant cell type correlations is distinct in each progression group, with 70 correlation pairs in slow progressors, 64 pairs in standard progressors, and 52 pairs in fast progressors, suggesting that highly coordinated immune responses present in slow progressors are absent, or break down, in more rapidly advancing disease. Functional categorization revealed that slow and standard progressors exhibit enhanced adaptive immune coordination, with 31% and 33% of their significant correlation pairs occurring between adaptive immune cells (vs. 22% in healthy controls), while showing reduced coordination between innate immune cells (16% vs. 24% in healthy). This shift toward enhanced adaptive immune coordination potentially represents compensatory mechanisms that dampen progression through better T and B cell coordination(Beers et al., 2017; Yazdani et al., 2022).

On the other hand, fast progressors maintain coordination proportions most similar to healthy controls yet exhibit the poorest outcomes. This suggests that the identity of coordinating cells and cell hubs, not just the overall balance between adaptive, innate and innate-adaptive coordination, determine functional immune outcomes. Second, fundamental cell hubs mediating crosstalk are indeed also distinct in disease and shift from B cell and basophil coordination in healthy controls to neutrophil and T cell-centered coordination across all ALS groups. Third, we observed striking differences in the direction of correlation among several NK cell mediated cell-cell relationships between progression groups suggesting that immune coordination patterns undergo fundamental reorganization despite being seemingly stable based on population frequency alone. Most strikingly, NK cells exhibit positive correlations with T helper cells in standard progressors (r=0.50) but show negative correlations with these cells in fast progressors (r=-0.87), despite their unchanged frequency across progression groups (Murdock et al., 2021; Piccoli et al., 2023). NK cells can directly regulate T-helper frequency and polarization through cytotoxicity and cytokine secretion, respectively (Pallmer & Oxenius, 2016).

It is therefore possible that NK cells differentially regulate ALS progression through direct interactions with CD4+ T-cell subtypes that confer protective (e.g. Th2) or maladaptive (e.g. Th1) states. Finally, seven correlation patterns are conserved across all ALS progression groups while absent in healthy controls, representing disease-associated immune signatures. The strongest of these is the negative correlation between neutrophils and T helper cells (r=-0.93), which validates the observation that neutrophil-to-lymphocyte ratios predict poor ALS survival (Choi et al., 2020; Wei et al., 2022).

Importantly, key cell populations identified by analysis of progression-specific frequency patterns proved even more informative through their relationships with other cell types. For example, coordination between pDCs and T-regs emerged as key discriminative features for distinguishing rapid from non-rapid progression, which directly connects our findings to existing therapeutic landscapes where T-reg-based cellular therapies are under clinical investigation (Alsuliman et al., 2016; Beers et al., 2022; Kim et al., 2018; Shneider et al., 2025; Thonhoff et al., 2018). Because pDCs directly drive T-reg expansion and maintenance(W. Chen et al., 2008), it is possible that the loss of pDCs is intimately linked with reduced T-reg frequencies due to the inability to incite tolerogenic adaptive immunity which in turn leads to a rapidly advancing disease-state. Additionally, our feature importance analysis revealed that T cell subset interactions, especially central and effector memory (CM/EM) CD4+ with T helper cells and naive CD4+ populations, are most discriminative for healthy versus ALS classification, suggesting these relationships hold potential to be targets of intervention. Altogether, these findings suggest that therapeutic interventions should prioritize restoring immune coordination patterns rather than simply targeting individual cell populations.

Notably, conserved immune coordination patterns across studies can be readily detected even with different experimental technologies, as evidenced by successful cross-platform model validation, which we suspect is related to biological relevance across modalities. For rare diseases like ALS, where patient cohorts are inherently limited, this cross-platform robustness enables training models on one platform and validating them on independent datasets from other platforms, significantly expanding the utility of existing datasets for biomarker discovery and validation.

Although our study revealed several robust immune signatures correlating with ALS progression, there are limitations to address in future efforts. Longitudinal studies tracking immune cell coordination dynamics are needed to determine whether the coordination patterns we observed are stable characteristics of different progression groups or change dynamically within individual patients. Such studies could potentially identify intervention windows when compensatory mechanisms remain active. Experimental validation through in vitro co-culture experiments or in vivo studies, which modulate one cell-type within a prominent coordinating pair can be implemented to observe the overall effect on inflammation. These insights can ultimately be harnessed to develop new therapeutic strategies to modulate disease progression (Schwartz & Colaiuta, 2024). Additionally, integration of functional immune markers could further enhance the predictive power of our models and provide mechanistic insights into why certain immune relationships are preserved or lost in different progression contexts. While our focus on peripheral immune signatures captures systemic immune dysfunction that is known to both reflect and contribute to ALS pathogenesis through bidirectional peripheral-CNS immune communication (Beers et al., 2017; Murdock et al., 2017; Yazdani et al., 2022), future studies should determine whether these coordination patterns can predict therapeutic responses.

In summary, we identified critical immune coordination patterns in blood, which can better stratify ALS progression groups over traditional single target biomarker approaches. Importantly, we demonstrate that these coordination patterns can be translated into predictive models that successfully classify disease onset and progression rates across independent cohorts and single-cell technologies. Our findings highlight several key immune components as primary drivers of disease progression and potential targets for future therapeutic strategies. The use of a coordinated network-based approach may extend beyond ALS to other neurodegenerative diseases characterized by immune dysfunction.

## Methods

### Study Design and Participants

In this study, we used a discovery-validation design where two independent cohorts were analyzed with different single-cell technologies. The discovery cohort consisted of 44 participants: 35 ALS patients and 9 healthy controls recruited from Duke University (Institutional Review Pro00109640) and UNC Chapel Hill (Institutional Review 22-2638) interdisciplinary ALS clinics following provision of written informed consent. ALS patients were classified into progression groups based on the rate of change in the revised ALS functional rating scale (ΔALSFRS-R/t scores): slow progression (n=16, ΔALSFRS-R/t < 0.5), standard progression (n=9, 0.5 ≤ ΔALSFRS-R/t ≤ 1.0), and fast progression (n=10, ΔALSFRS-R/t > 1.0). Patient demographics showed similar sex distributions across groups, with slightly younger age of onset in fast progressors (59.4 ± 10.1 years) compared to slow (65.6 ± 11.6) and standard (63.9 ± 7.5) progression groups.

The validation cohort consisted of 40 participants from the publicly available dataset by Itou et al. (2024) obtained from NCBI GEO (accession# GSE244263): 30 ALS patients and 10 healthy controls analyzed via single-cell RNA sequencing of peripheral blood mononuclear cells. ALS patients in the validation cohort were classified as either Rapid progressors (ΔALSFRS-R/t ≥ 1.0, n=7) or Non-Rapid progressors (ΔALSFRS-R/t < 1.0, n=23). To align with this classification scheme, we mapped our discovery cohort’s slow and standard groups to Non-Rapid, and our fast group to Rapid for validation analyses.

#### ALSFRS-R/t Score Calculation

The rate of disease progression was quantified using the ΔALSFRS-R/t score (Cedarbaum et al., 1999), calculated as:

ΔALSFRS-R/t = (48 - ALSFRS-R_current) / (time_current - time_onset), where 48 represents the maximum possible ALSFRS-R score, ALSFRS-R_current is the patient’s score at time of sample collection, time_current is the time of assessment, and time_onset is the time of symptom onset. This metric provides a standardized measure of functional decline rate, with higher values indicating faster disease progression.

### CyTOF Data Acquisition and Preprocessing

#### Sample Collection and Processing

Patient blood was collected at UNC Chapel Hill and Duke University Neurology clinics following provision of informed consent. Blood was collected in standard 5.0 mL heparinized vacutainer collection tubes (Becton Dickinson). Approximately 300 μL of anticoagulated whole blood was added to the lyophilized antibody pellet of a Maxpar Direct Immune Profiling Assay (MDIPA) tube (Standard BioTools Inc.) and allowed to incubate for 30 minutes at room temperature. To ensure even staining, the tube was gently agitated every 10 minutes. Next, stained blood was added to 420 μL of Smart Tube Proteomic Stabilizing agent (Smart Tube Inc.) and allowed to incubate for 10 minutes at room temperature before immediate storage at-80°C where they awaited batch processing. Samples were collected from August 2022 through November 2024 and analyzed in three batches. One day prior to analysis, batches of stained blood samples were thawed and transferred to 10 mL of 1X Thaw/Lyse buffer (Smart Tube Inc.). Samples were incubated at room temperature for 10 minutes with orbital shaking, followed by centrifugation at 500X RCF for 7 minutes. This lysis and centrifugation step was repeated once more. Next, the cell pellets were resuspended in 10 mL of 1X Cell Staining Buffer (Standard BioTools Inc.) and centrifuged as above. The supernatant was removed and the cell pellets were resuspended in 1 mL of Maxpar phosphate buffered saline (Standard BioTools). The cell resuspension was then filtered through a 70 μm filter directly into 1 mL of fresh 4% paraformaldehyde (Electron Microscopy Services) for a final concentration of 2% paraformaldehyde. Samples were then incubated at 4°C for 1 hour. Finally, cells were collected by centrifugation at 800X RCF for 7 minutes and resuspended in 1 mL of Maxpar Fix/Perm Buffer with 1:3000 Cell-ID Iridium Intercalator (Standard BioTools) and incubated at 4°C for 24 hours. Prior to sample acquisition, cells were collected by centrifugation, washed sequentially in Cell Staining Buffer and Cell Acquisition Solution (CAS, Standard BioTools) and resuspended inCAS containing 10% EQ Four Element calibration beads (Standard BioTools) at 600,000 cells/mL. Samples were analyzed on a Helios Mass Cytometer equipped with Fluidigm software (v7.0) and a minimum of 150,000 events were collected per sample. FCS Data files (flow cytometry standard) were then cleaned of beads, debris, doublets, and dead cells using CytoBank (v10.7).

#### Data Preprocessing and Quality Control

CyTOF data stored in FCS files were processed using FlowKit to parse individual events and their respective information across channels from three acquisition batches. Quality control filtering was implemented through dual-threshold filtering on DNA intercalation channels (191Ir and 193Ir). DNA signal was calculated as the sum of both iridium channels, and filtering thresholds were established using percentile-based cutoffs so that the 1st percentile served as the lower threshold to remove cellular debris, while the 99th percentile served as the upper threshold to remove cell aggregates and doublets, respectively. Cells with DNA signal between these thresholds were retained for downstream analysis.

A panel of 30 markers was retained for analysis, including lineage markers (CD3, CD4, CD8a, CD19, CD20, CD14, CD16, CD56), activation and differentiation markers (CD25, CD27, CD28, CD38, CD45RA, CD45RO, CD57, HLA-DR), chemokine receptors (CCR4, CCR6, CCR7, CXCR3, CXCR5), and specialized markers (CD123, CD127, CD161, CD183, CD185, CD194, CD196, CD294, TCRγδ, CD66b, IgD). The expression of each marker was transformed using the inverse hyperbolic sine (arcsinh) function with a cofactor of 5 as f(x) = arcsinh(x/5) = ln(x/5 + √((x/5)² + 1)).

This transformation stabilizes variance across the dynamic range and normalizes expression distributions for downstream analysis.

To ensure balanced representation across samples and prevent bias arising from variation in the number of profiled cells, we implemented balanced downsampling to exactly 50,000 cells per sample. There were variable numbers of cells across samples, ranging from 80,828 to 783,269 cells per sample. After quality control filtering and balanced sampling, the final dataset contained 2,200,000 cells across 44 samples for subsequent analysis.

All samples were processed and combined into a single AnnData object (using the anndata package in Python) with batch identity, sample information including age and sex, patient IDs, and health status preserved as metadata for subsequent analysis steps.

#### Cell Type Identification and Annotation

Following data preprocessing and balanced sampling, batch effect correction was performed using ComBat (Johnson et al., 2007; Leek et al., 2017; Pedersen, 2012/2025) as implemented in scanpy to account for technical variation across the three acquisition batches. After batch correction, the values from each channel were scaled to have a maximum threshold of 10 to prevent outliers from dominating the analysis while maintaining relative marker expression differences.

Dimensionality reduction was applied to cells by performing Principal Component Analysis (PCA) with 40 components, followed by neighborhood graph construction using *k =* 5 nearest neighbors. Cells were clustered into cell types using the Leiden algorithm with a resolution of 1.5 to achieve fine-grained separation of immune cell populations (Traag et al., 2019). UMAP embedding was computed for visualization purposes. This unsupervised clustering approach yielded 64 distinct cell clusters from the 2,200,000 cells.

Cell types were annotated according to the z-score normalized marker expression patterns across all 30 markers (Fig. 2). Cell types were ultimately identified based on the Maxpar Direct Immune Profiling Assay panel guidelines (Standard BioTools, 2022). Our annotation identified major immune cell lineages including T cells (naive and memory CD4+ and CD8+ subsets, regulatory T cells, T helper cells, and γδ T cells), B cells (naive, memory, transitional, and plasmablasts), NK cells, monocytes (classical, non-classical, and transitional), plasmacytoid dendritic cells, granulocytes (neutrophils, eosinophils, basophils), and innate lymphoid cell precursors. This manual annotation process consolidated the 64 initial clusters into 25 biologically meaningful cell types. Cells with low overall marker expression were excluded from the analysis.

### Statistical Analysis of Cell Type Frequency

In each sample, the frequency for each cell type was computed as the proportion of that sample’s cells assigned to the given cell type. The computed frequencies of all cell types were log-transformed for subsequent analysis. Statistical tests to evaluate frequency differences in individual cell types between progression groups were implemented using non-parametric tests. Granulocytes (neutrophils, basophils, and eosinophils) were excluded from frequency difference analyses because their high abundance in whole blood (neutrophils comprised 66.6% of all cells) results in very small frequencies for other immune cell populations, limiting the statistical power to detect meaningful differences in non-granulocyte populations. For overall group comparisons across the four groups (healthy, slow, standard, fast), we applied Kruskal-Wallis tests. Pairwise comparisons between specific groups were conducted using Mann-Whitney U tests. Multiple testing correction was applied using the Benjamini-Hochberg procedure, with correction applied separately for Kruskal-Wallis tests across all cell types and within each cell type for Mann-Whitney U tests.

### Correlation Analysis of Cell Type Frequencies with Disease Progression

To assess relationships between frequencies of immune cell types and disease progression rates, we calculated Spearman’s rank correlations between log-transformed cell type frequencies and ΔALSFRS-R/t scores. Two different variants of correlations were computed. First, we did so, including healthy controls (assigned ΔALSFRS-R/t = 0) to assess relationships across the entire health-disease spectrum. Second, we excluded healthy controls to focus specifically on progression rate differences within ALS patients. The p-values of the computed correlations were corrected using the Benjamini-Hochberg procedure.

### Inter-Cell Type Frequency Correlation Analysis

To investigate coordination patterns between components of the immune system across disease states, we performed a comprehensive cell type-cell type frequency correlation analysis within each progression group. This analysis was motivated by our hypothesis that ALS progression rate is likely to be mediated by particular cellular coordination patterns, rather than simple changes in individual cell frequencies.

To address sample size imbalances between groups, we implemented balanced sampling by randomly selecting 9 samples from each group (matching the smallest group size) and repeated this process over 100 iterations to obtain robust estimations of the correlation between cell types based on their frequency patterns. For each iteration, cell type frequencies were calculated as proportions within each sample, with a small epsilon value (1×10⁻^¹⁰^) added to denominators to prevent division by zero. Frequencies were log(1+x) transformed to improve normality and reduce the influence of outliers. All pairwise Spearman correlations between cell type frequencies were computed within each progression group.

To ensure correlation stability, we implemented bootstrap resampling (n=1000 iterations) for each cell type pair, adding small amounts of Gaussian noise (σ=0.01) to the log-transformed frequencies to prevent identical values from artificially inflating correlation estimates.

Correlations with a coefficient of variation >200 for either cell type were excluded from analysis to remove unreliable estimates from rare cell populations.

Statistical significance of correlations between cell types based on their frequencies was assessed using Fisher’s z-transformation for meta-analysis across iterations. For each cell type pair, correlation coefficients were transformed using the inverse hyperbolic tangent (Fisher’s z), weighted by sample size (n-3), and combined across iterations. The combined z-statistic was calculated as,

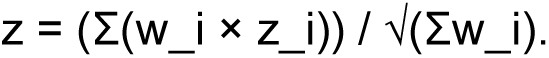

Here, w_i represents the weight (sample size - 3) and z_i the Fisher z-transformed correlation for iteration i. P-values were derived from the standard normal distribution of the combined z-statistic.

Final correlation estimates were obtained by averaging correlations across all 100 iterations, with correlation variance calculated as the variance of correlation coefficients across iterations. Multiple testing correction was applied using the Benjamini-Hochberg false discovery rate procedure within each progression group.

We applied stringent filtering criteria to identify reliable correlations: absolute correlation coefficient ≥ 0.5, FDR-corrected p-value < 0.001, and correlation variance ≤ 0.4. Coordination analysis was performed by representing cell types as nodes and significant correlations as edges, with node degree calculated to identify hub cells and correlation sign reversals systematically identified to detect fundamental changes in immune coordination patterns.

### Feature Engineering and Machine Learning to Predict ALS Onset and Progression

Building on our correlation coordination analysis that revealed progression-specific immune coordination patterns, we developed a systematic feature engineering approach to test the hypothesis that features that reflect coordination between immune cell types create more informative biomarkers than cell type frequencies alone. Our features were organized into distinct categories designed to capture distinct aspects of immune dysregulation.

Feature Categories and Biological Rationale

#### Basic Frequency Features

Log-transformed frequencies of all immune cell populations (Feature Set 1) or only significant cell types showing differences between ALS progression groups identified in the discovery cohort (Feature Set 2).

#### Relationship-Based Features

These features capture immune coordination patterns identified in our correlation analysis:

- **Log Ratio Features**: Calculated as the difference between log-transformed frequencies of two cell types: LogRatio(cell_i, cell_j) = log(freq_i + ε) - log(freq_j + ε) where freq_i and freq_j represent the frequencies of cell types i and j respectively, and ε = 1×10⁻⁶ was added to prevent logarithm of zero. These features capture the relationships between the abundances of different immune cell types.
- **Interaction Features**: Calculated as the product of cell type frequencies to model potential coordinated responses. Here, an interaction between a pair of cell types *i* and *j* can be stated as Interaction(cell_i, cell_j) = freq_i × freq_j. These features represent

potential synergistic relationships between cell populations, amplifying signals when both cell types change together.

#### Task-Specific Features

Features derived from correlations specifically relevant to each classification task, as identified in the discovery cohort. This approach focuses on the most relevant immune relationships for each classification challenge.

#### Combined Features

Integration of log-transformed individual cell frequencies with derived relationship features in various configurations, testing whether combining individual cell type frequency information with pairwise relationships enhances classification performance.

All cell type frequencies were log-transformed before feature engineering to improve distribution normality and model performance. The complete list of 21 engineered feature sets is provided in Supplementary Table S1.

### Machine Learning Classification

Random Forest classifiers were implemented for binary classification tasks using scikit-learn. Primary classification tasks included distinguishing Healthy from ALS patients and separating Non-Rapid from Rapid progressors. Random Forest was selected for its ability to capture non-linear relationships between immune features and robustness to overfitting on small datasets.

Model training employed a cross-platform validation approach where models were trained exclusively on CYTOF data and validated on scRNA-seq data to assess generalizability across platforms. To ensure robust performance estimates, the complete training and validation pipeline was repeated 5 independent times, generating multiple independent model instances for each feature set configuration. Final performance metrics represent the mean across these 5 runs.

Hyperparameter optimization was performed for each feature set using grid search with 3-fold cross-validation. To address inherent class imbalance, balanced down-sampling was applied within each of the 3 cross-validation folds to the training portion before model fitting, followed by an additional balanced down-sampling step during final model training (4 total balanced sampling steps per model). Each balanced down-sampling reduced each class to match the size of the smallest class, ensuring equal representation across progression groups. Optimized parameters included the number of estimators (100, 200, 500), maximum depth (None, 10, 20), minimum samples split (2, 5), and minimum samples per leaf node (1, 2). The 3-fold approach ensured sufficient samples per fold given the size of our discovery cohort.

Model performance was assessed using 200 iterations of bootstrap resampling on the scRNA-seq validation cohort to generate robust confidence intervals for multiple complementary metrics appropriate for imbalanced datasets: accuracy for overall classification performance, area under the ROC curve (AUC) for discrimination ability across all decision thresholds, average precision for performance assessment on imbalanced datasets, and F1 score as the harmonic mean of precision and recall.

Feature importance was calculated using built-in Random Forest importance scores based on mean decrease in impurity.

### scRNA-seq Validation Cohort Preprocessing

#### Data Preprocessing

The validation cohort consisted of publicly available single-cell RNA sequencing data from Itou *et al*. (2024) obtained from NCBI GEO (accession# GSE244263). Raw count matrices were processed using scanpy, following established single-cell RNA seq best practices to ensure compatibility with CyTOF-derived features for cross-platform validation as described below.

Individual sample files were processed with balanced sampling (maximum 750 cells per sample) to manage computational load while maintaining representative populations. Quality control metrics were calculated using scanpy, followed by normalization to 10,000 counts per cell and log(1+x) transformation of all measured transcripts. Highly variable genes were identified using the Seurat method, and only the 3000 most variable genes were retained for downstream analysis to reduce computation burden and focus on the most informative transcriptional features.

Batch effect correction was performed using Harmony (Korsunsky et al., 2019) on the 3,000 most variable genes to account for technical variation across samples while preserving biological differences between disease states. Each cell was represented by its 35 principal components. A graph representation of cells was created for subsequent clustering and UMAP visualization by connecting each cell with its 15 nearest neighbors.

Cell clustering was performed using the Leiden algorithm with resolution=1.5, generating 34 distinct clusters for manual annotation. Cell type annotation was conducted through systematic examination of canonical immune cell marker expression patterns, establishing cell type definitions based on established lineage markers and functional states (Supplementary Figure S4). The annotation strategy identified major immune cell populations including T cell subsets (CD4⁺ and CD8⁺ naive, memory, effector, and regulatory populations), B cell populations (naive, memory, transitional, and plasma cells), monocyte subsets (classical, non-classical, transitional), NK cells, dendritic cells, neutrophils, and other myeloid populations.

#### Cross-Platform Model Validation

Models were trained exclusively on CyTOF-derived features from the discovery cohort and applied to corresponding features derived from the scRNA-seq validation cohort. Feature compatibility was ensured by mapping cell type frequencies between platforms based on manual annotation consensus. This external validation approach provided an assessment of model generalizability and biological relevance of identified immune signatures across different analytical platforms and sample types.

Prediction accuracy in the validation cohort was assessed using accuracy, AUC, average precision, and F1 score. Statistical significance of performance differences between the best-performing feature sets and the baseline all cell types feature set was assessed using Welch’s t-test on validation cohort predictions with significance set at α = 0.05.

### Statistical Software and Implementation

All analyses were performed in Python 3.8+ using established scientific computing packages. Single-cell analysis workflows used scanpy version 1.9.0+, machine learning implementations used scikit-learn version 1.3.0+, and data manipulation employed pandas version 2.0.0+ and numpy version 1.24.0+. Statistical tests were conducted using scipy version 1.10.0+, and visualization used seaborn and matplotlib. Batch correction employed ComBat implementation in scanpy for CyTOF data (Johnson et al., 2007; Leek et al., 2017; Pedersen, 2012/2025), and Harmony integration for scRNA-seq data (Korsunsky et al., 2019).

Statistical significance was set at α = 0.05 for all tests, with multiple testing corrections applied as specified for each analysis type. For cell-cell frequency correlation analysis, we used a more stringent significance threshold of α = 0.001 to identify the most robust immune coordination patterns.

## Data Availability

The single-cell RNA sequencing data (‘Validation cohort’) are publicly available from Itou et al. (2024) and can be accessed at NCBI GEO (accession# GSE244263). Processed AnnData objects for both the mass cytometry data (‘Discovery cohort’) and single-cell RNA sequencing data (‘Validation cohort’) have been deposited in Zenodo (DOI: 10.5281/zenodo.16039496). Code for the machine learning models are available on GitHub at https://github.com/ldhawka/als-cross-platform-classification.

## Contributions

L.D. and B.A.E conceptualized study, designed, and performed experimentation. O.K.A., K.P., and R.B.M assisted with blood processing protocols. M.A.I. performed mass cytometry. R.T., X.L., and R.B., provided patient care for study participants, managed clinical data, and facilitated blood sample procurement. T.J.C. and N.S. supervised generation, acquisition, and analysis of research data. L.D., B.A.E, T.J.C., and N.S. wrote the original manuscript, which was reviewed and edited by all authors.

## Conflict of interest

The authors declare no competing conflict of interest.

## Study Approval

The study was authorized by Institutional Review Pro00109640 of Duke University and 22-2638 of University of North Carolina at Chapel Hill. All participants were studied after provision of written informed consent.

## Acknowledgments

UNC Mass Cytometry Core is supported by the University Cancer Research Fund (UCRF) UNC Cancer Center Core Support Grant P30CA016086. B.A.E was supported in part by the National Institute of Neurological Disorders and Stroke F31 NS122242. Clinical work was supported in part by the North Carolina Translational and Clinical Sciences (TraCS) pilot award 550KR282107 awarded to R.T. and X.L.. T.J.C. was supported in part by R21AG084251 awarded by the National Institute on Aging. N.S. was supported in part by 5R21AI171745-02 awarded by the National Institute of Allergy and Infectious Diseases of the NIH. We’d like to thank C. Simmons, M. Chopra, M. Ward, and H. Zampa for patient recruitment, enrollment, and sample acquisition. Finally, we sincerely thank the ALS patients and family members for their contribution to this work.

## Notes

https://github.com/ldhawka/als-cross-platform-classification

https://docs.google.com/spreadsheets/d/1k2xTGeLSk7UaLs_5XpgnEo-xw5kHLB7_9D-5whFw__Q/edit?usp=sharing

### Competing Interest Statement

The authors have declared no competing interest.

