## Supplementary figures and tables for "Peripheral immune patterns enable robust cross-platform prediction of ALS onset and progression"

**Supplementary Table S1**

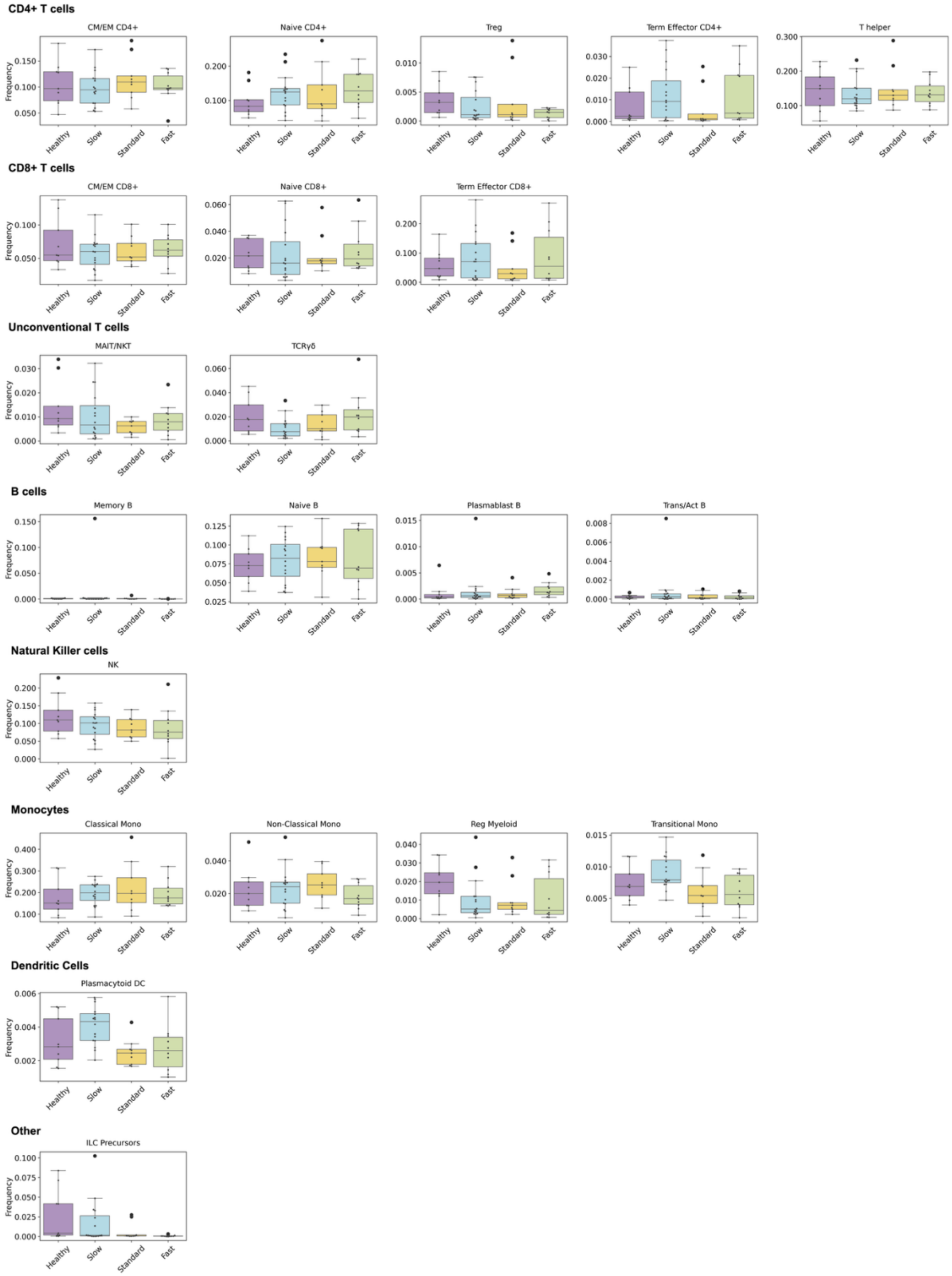


Fig 1: Cell type frequency changes across ALS progression groups across major immune

lineages, excluding granulocytes.

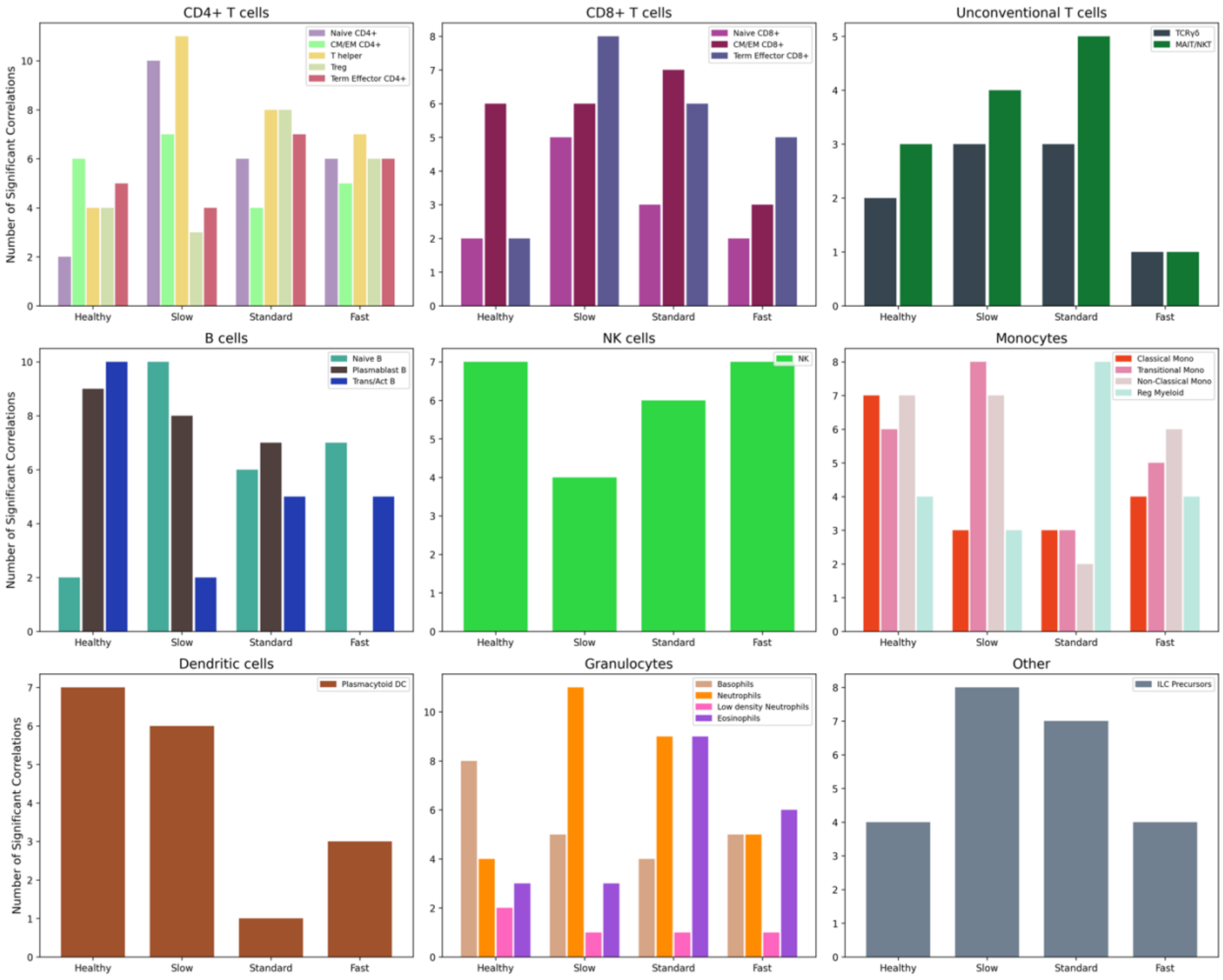


Fig 2: Node degree of each cell type reflecting the overall strength of statistically significant correlations with other cell types in each progression groups

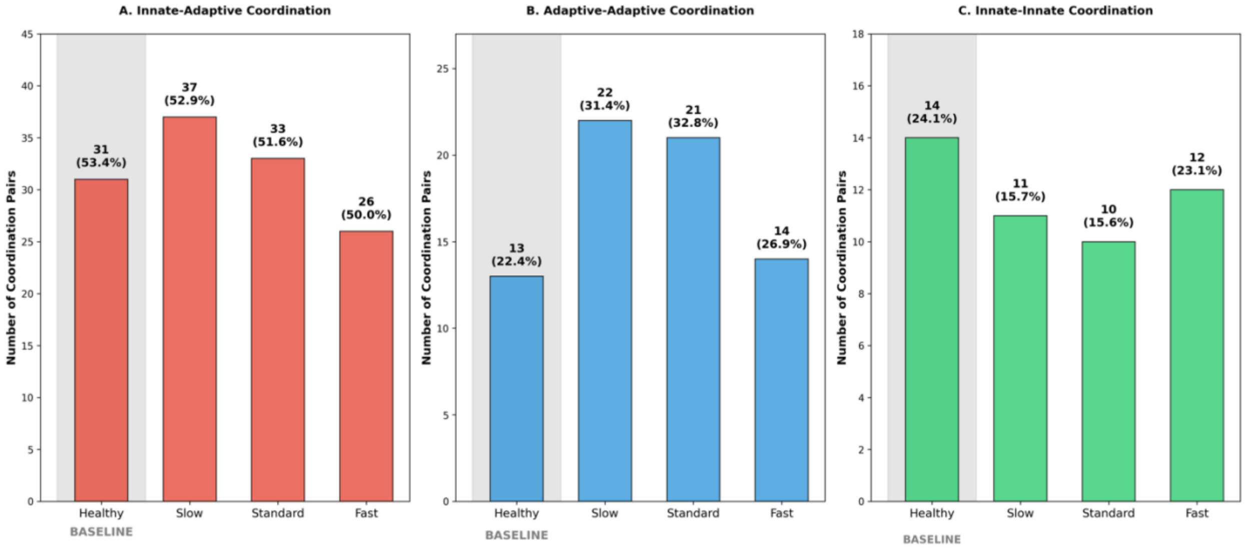


Fig 3: Immune coordination patterns reveal distinct signatures across ALS progression groups. Significant correlation pairs ($|r| \geq 0.5$, FDR $p < 0.001$) were categorized by immunity type. (A) Innate-adaptive coordination. (B) Adaptive-adaptive coordination. (C) Innate-innate coordination. Numbers show counts and percentages of total pairs. Gray shading indicates healthy baseline. Slow and Standard progressors show enhanced adaptive-adaptive coordination (31-33% vs 22% healthy) with reduced innate-innate coordination (16% vs 24% healthy), while Fast progressors maintain healthy-like patterns across all types.

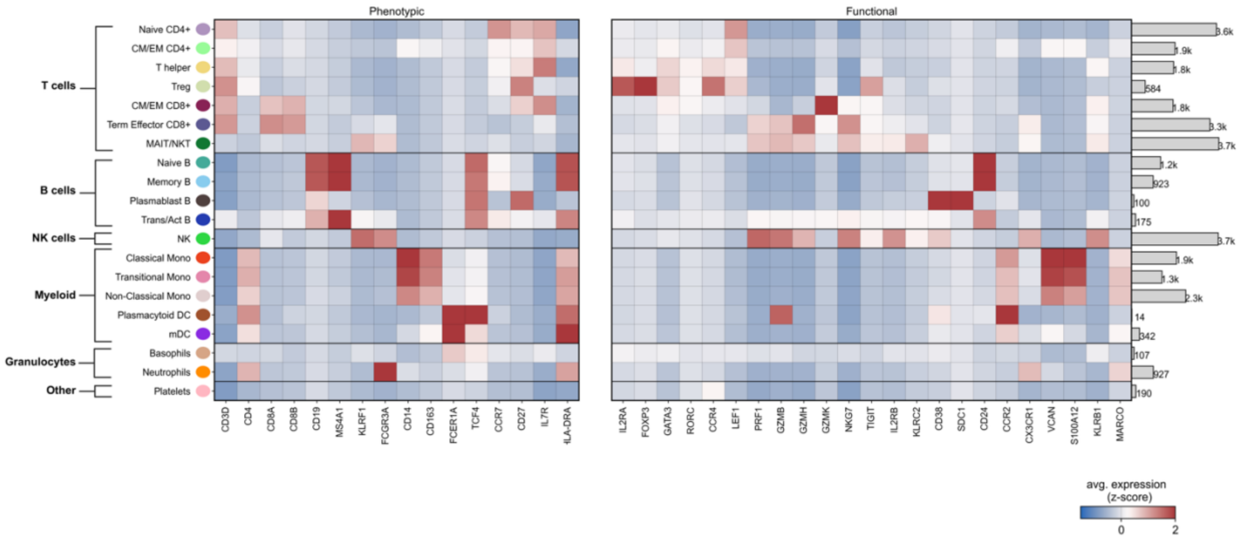


Fig 4: Heatmap showing average gene expression (z-score) of phenotypic (left) and functional (right) markers used to phenotype cell type populations in the scRNA seq validation cohort from Itou et al. 2024. The number on the right of the heatmap indicates the frequency counts of each cell type across all samples. Cell types are grouped by major lineages (T cells, B cells, NK cells, Myeloid, Granulocytes, and Other).
